# Bmf Facilitates Protein Degradation and Reduces Beclin1 Ubiquitination to Inhibit Autophagy Independent of mTOR

**DOI:** 10.1101/2020.01.02.892828

**Authors:** Monica Delgado-Vergas, Susan Fort, Dereje Tassew, Yohannes Tesfaigzi

## Abstract

Previous observations suggested that Bcl-2 modifying factor (Bmf) affects autophagy but the underlying mechanisms were unknown. The present studies show that Bmf inhibited the initiation and flux of autophagy in a manner that is independent of the mTOR pathway and inhibition of mTOR increased Bmf expression to temper the autophagic cell death. In mice, emphysema was observed in Bmf-deficient mice, suggesting that Bmf suppresses autophagic cell death of alveolar type II cells. Bmf deficiency increased ubiquitination of Beclin1 with K63 chains and released Beclin1 from Bcl-2. However, Bmf deficiency also increased the levels of polyubiquitinated proteins in general. In mice, Bmf-deficiency robustly increased p62 levels in all tissues analyzed, but LC3-II levels were reduced only in the hearts of old mice. Also, Bmf-deficiency caused persistent mucous cell metaplasia in mice exposed to allergen and increased levels of polyubiquitinated Muc5ac in differentiated airway epithelial cells. The reduction of ubiquitinated proteins was mediated by the BH3- and dynein binding-domains of Bmf. Together, these findings show that the primary role of Bmf is to reduce protein levels and affects K63- and K48-ubiqutination.

## Introduction

The Bcl-2 family of proteins regulates the intrinsic apoptotic pathway by integrating signals that originate from a variety of cellular stressors. The Bcl-2 subfamily, known as the BCL-2 homology domain 3 only containing (BH3-only) proteins (Bim, Puma, Bid, Bad, Bmf, Bik, Hrk and Noxa) initiate apoptosis in response to DNA damage (Villunger, Michalak et al., 2003) or cytokines (Ekoff, Kaufmann et al., 2007, Mebratu, Dickey et al., 2008). They either bind to neutralize pro-survival Bcl-2 family members or relieve the inhibition of pro-apoptotic Bax and Bak.(Happo, Strasser et al., 2012) While their pro-apoptotic function has been studied well, few studies have focused on the non-cell death-related roles of the Bcl-2 family of proteins. Bcl-2 is implicated in regulating cell cycle entry (Huang, O’Reilly et al., 1997, Linette, Li et al., 1996), and in inhibiting autophagy by binding Beclin1 (Fernandez, Sebti et al., 2018, Pattingre, Tassa et al., 2005). Bad affects insulin secretion and metabolism (Danial, Walensky et al., 2008), Noxa by cross-linking to phospho-HSP27 stabilizes I□B□ and regulates inflammation (Zhang, Jones et al., 2018), while Bid by interacting with NOD1, NOD2, and the IkB kinase (IKK) complex facilitates NF-kB activation.(Yeretssian, Correa et al., 2011)

Loss of Bcl-2 modifying factor (Bmf) protects lymphocytes against apoptosis induced by glucocorticoids or histone deacetylase inhibition.(Pfeiffer, Halang et al., 2015) Moreover, *bmf^-/-^* mice develop a B cell-restricted lymphadenopathy caused by the abnormal resistance of these cells to a range of apoptotic stimuli. *Bmf* mRNA is increased upon loss of matrix attachment or disruption of the actin cytoskeleton and Bmf initiates anoikis, while down-regulation of Bmf expression by small interfering RNAs is sufficient to prevent anoikis (Schmelzle, Mailleux et al., 2007). Finally, Bmf-deficiency accelerates the development of gamma irradiation-induced thymic lymphomas, suggesting that Bmf plays a role in apoptosis signaling and can function as a tumor suppressor.(Labi, Erlacher et al., 2008)

At baseline, Bmf mediates fetal oocyte loss in mice at E15.5 and PN1 Primordial follicle numbers remain elevated throughout reproductive life and confer prolonged fertility in *bmf^-/-^* females (Liew, Nguyen et al., 2017, Liew, Vaithiyanathan et al., 2014) However, the major phenotype of *bmf^-/-^* mice is a defect in utero vaginal development, including an imperforate vagina and hydrometrocolpos (Hubner, Cavanagh-Kyros et al., 2009). Because vaginal introitus formation requires the apoptosis of the vaginal mucosa (Rodriguez, Araki et al., 1997), it is assumed that Bmf facilitates this formation by apoptotic mechanisms. However, cytochrome c pathway does not fully account for the pro-apoptotic actions of Bmf because *Apaf1^-/-^* (Yoshida, Kong et al., 1998) or *Casp9^-/-^* (Hakem, Hakem et al., 1998, Kuida, Haydar et al., 1998) mice do not show this phenotype. Therefore, it is not clear whether Bmf deficiency causes the phenotype due to its apoptotic function or by another mechanism.

We reported that deacetylation of p53, which activates nuclear p53, suppresses Bmf expression. This loss of Bmf diminishes the interaction of Beclin1 and Bcl-2 and thereby facilitates autophagosome formation.(Contreras, Mebratu et al., 2013) However, how this BH3-only protein, rather than competing for, enhances the BH3-mediated interaction of Beclin-1/Bcl-2 to inhibit autophagy was not clear.

Autophagy is a conserved cellular degradation pathway to digest intracellular components during starvation or related stress to provide a source of amino acids for the synthesis of new proteins.(Takeshige, Baba et al., 1992) Phosphorylation of mammalian target of rapamycin (mTOR) inhibits autophagy by controlling UNC-51-like kinase 1 (ULK1) ubiquitination (Nazio, Strappazzon et al., 2013). The activated ULK1/2 kinase complex, including focal adhesion kinase family interacting protein of 200 kDa (FIP200), and subsequent activation of the Beclin-1–Vps34–AMBRA1 complex are necessary to initiate phagophore formation (Nazio et al., 2013). Among others, p62 recognizes K63-polyubiquitinated proteins and phase separates proteins into larger condensates to package them into the autophagosomal membrane. (Danieli & Martens, 2018, Filimonenko, Isakson et al., 2010) Microtubule-associated protein light chain 3 (LC3) I is proteolytically cleaved and attached to phosphatidylethanolamine to form a lipidated LC3-II and facilitate autophagosome maturation. Therefore, conversion of LC3B-I to LC3B-II indicates the presence of mature autophagosomes (Moreau, Ravikumar et al., 2011). Finally, lysosomes fuse with autophagosomes, and the resulting autophagolysosomes and their contents are degraded.

The current study was focused on clarifying the molecular mechanisms how Bmf regulates autophagy. We found that Bmf is a regulator of proteasomal degradation, and that deficiency in Bmf causes accumulation of protein aggregates. Bmf deficiency facilitates Beclin1 ubiquitination by the K63 polyubiquitin chain to induce autophagy and cause the death of alveolar epithelial cells to cause emphysema in mice. Among other proteins, following injury to the lung, Bmf may also facilitate resolution of mucins in airway epithelia likely by driving their degradation.

## Results

### Bmf inhibits autophagic initiation and flux

We have previously shown that mouse embryonic fibroblasts (MEFs) or mouse airway epithelial cells (MAECs) from *bmf^-/-^* compared with *bmf^+/+^* mice present with more autophagosomes (Contreras et al., 2013). To investigate whether the accumulated autophagosomes represent a real induction of autophagy or just inhibition of the autophagic flux, we treated cells with bafilomycin A1 (BAF), an inhibitor of the lysosomal degradation. Increased LC3-II levels were observed in vehicle-treated *bmf^-/-^* compared with *bmf^+/+^* MEFs and this difference was enhanced by BAF (**Figure 1A**). Similar results were observed in *bmf^+/-^* compared with *bmf^+/+^* MEFs after BAF treatment (data not shown), supporting the idea that both the initiation and flux of autophagy were increased in Bmf-deficient cells. In contrast to MEFs, in *bmf^-/-^* (knockout) compared with *bmf^+/+^* MAECs, the difference in LC3-II levels was minimal, even in the presence of BAF (**Figure 1B, left blot**). However, LC3-II accumulation was more evident in heterozygous (*bmf^+/-^*) MAECs, especially after BAF treatment (**Figure 1B, right blot**). Western blot analysis confirmed that Bmf protein is present in *bmf^+/+^* and *bmf^+/-^* but not in *bmf^-/-^* MAECs (**Figure S1A**), although >80 μg of protein extract was needed to detect Bmf protein. Further, reduced *bmf* mRNA levels in *bmf^+/-^* MAECs was confirmed by qPCR (**Figure S1B).** These results support our previous report that induction of autophagy by IFN-γ is mediated by reduced Bmf levels (Contreras et al., 2013).

**Figure 1:**
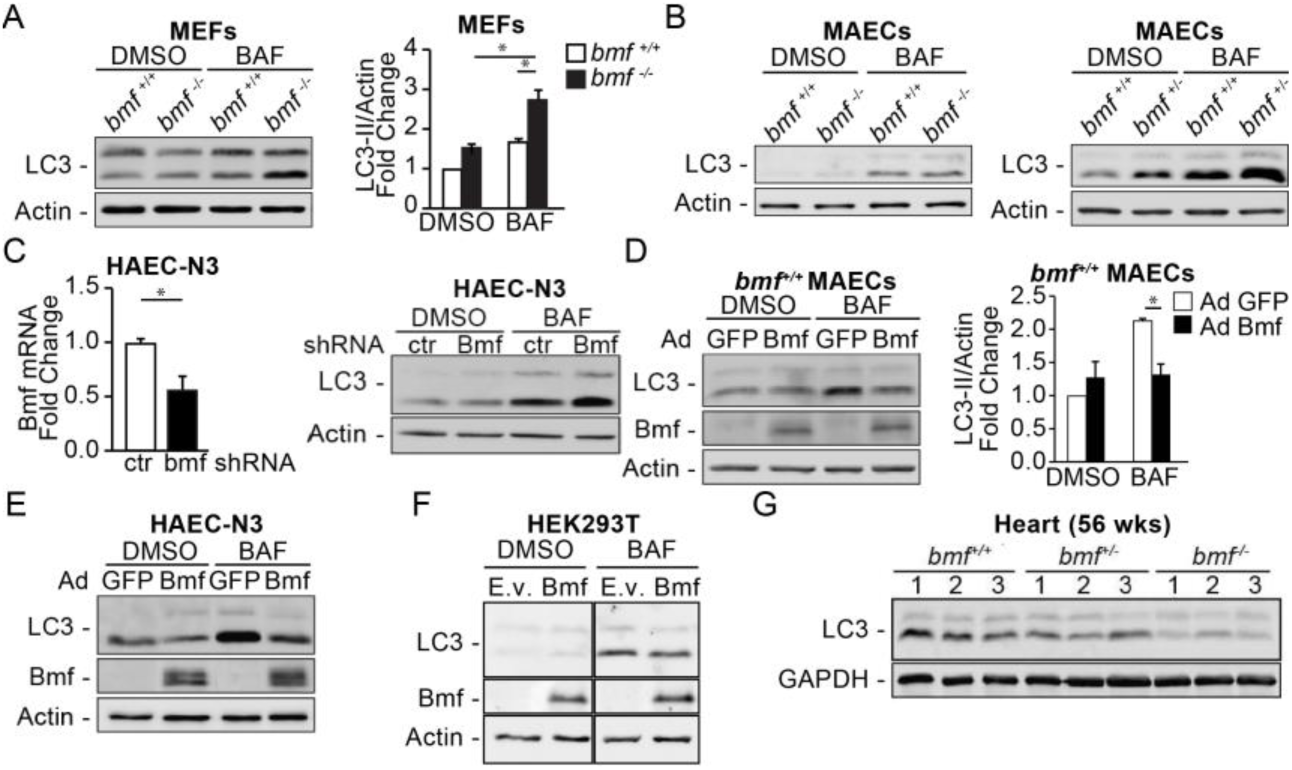
Bmf inhibits autophagic initiation and flux. **A)** Western blot analysis of protein extracts from *bmf^+/+^* and *bmf^-/-^* MEFs treated with DMSO or BAF (50 nM) for 1 h (left) and graph showing fold change of LC3-II/Actin (n= at least 3 independent experiments) (right); ANOVA, * P<0.05. **B)** Western blot analysis of protein extracts from *bmf^+/+^* and *bmf^-/-^* MAECs (left) and *bmf^+/+^* and *bmf^+/-^* MAECs (right) treated with DMSO or BAF (100 nM) for 4 h. **C)** Quantification of Bmf mRNA from HAEC-N3 cells infected with retroviral constructs with shRNA control (ctr) or sh-Bmf (n= at least 3 independent experiments) and Western blot analysis of same cells treated with DMSO or BAF (100 nM) for 4 h (right); T-test, * P<0.05. **D)** Western blot analysis of protein extracts from *bmf^+/+^* MAECs infected with AdGFP or AdBmf (MOI: 50) for 24 h and treated with DMSO or BAF (100 nM) for the last hour (left) and graph showing fold change of LC3-II/Actin (n= at least 3 independent experiments) (right); ANOVA, * P<0.05. **E)** Western blot anlaysis of protein extracts from HAEC-N3 cells infected with AdGFP or AdBmf (MOI: 50) for 24 h and treated with DMSO or BAF (100 nM) for the last 4 h. **F)** Western blot analysis of protein extracts from HEK293T cells transfected with empty vector (E.v.) or Bmf_S_ expressing plasmid for 24 h and treated with DMSO or BAF (100 nM) for the last 4 h. **G)** Western blot analysis of protein extracts from the heart lysates (100 µg or proteins) from naive 56 wks old male *bmf^+/+^*, *bmf^-/-^*, and *bmf^-/-^* mice.

To confirm these findings from murine MEF and MAECs in human airway epithelial cells (HAECs), Bmf expression was suppressed in the HAEC-N3 cells using a lentiviral vector expressing bmf shRNA. Because good antibodies to detect the human Bmf protein are unavailable, we confirmed the bmf knock down by analyzing the bmf mRNA levels by qPCR (**Figure 1C**). While LC3-II levels are similar in shBmf and shControl (shCtr) cells in the presence of DMSO, BAF treatment caused a greater accumulation of LC3-II in shBmf compared to control cells (**Figure 1C**). Reducing Bmf levels increased autophagy also in two other non-transformed HAEC lines, HAEC-N1 (**Figure S1C**) and AALEB cells (**Figure S1D**).

In contrast, adenoviral expression of Bmf (AdBmf) compared with GFP (AdGFP) in *bmf^+/+^* MAECs reduced accumulation of LC3-II when the autophagic flux was inhibited with BAF although similar levels of LC3-II were observed in the presence of DMSO only (**Figure 1D**). Similar results were observed in *bmf^+/-^* MAECs when infected with AdBmf compared with AdGFP (**Figure S1E**). HAEC-N3 cells also showed reduced LC3-II accumulation in AdBmf-compared with AdGFP-infected controls before and after BAF treatment (**Figure 1E**) as did HAEC-N1 cells (**Figure S1F**). Overexpression of Bmf by transfecting HEK293T cells with a plasmid expressing Bmf_S_, also reduced accumulation of LC3-II in the presence of BAF (**Figure 1F**). These observations show that Bmf inhibits autophagy also in human epithelial cells.

While autophagy is detected in Bmf-deficient cells in culture, the lung and thymus from *bmf^-/-^* compared with *bmf^+/+^* mice do not display difference in autophagy markers (Contreras et al., 2013). Even 24 h after injecting mice with BAF (2 µg per mice) to block the autophagic flux, LC3-II levels were not increased in *bmf^+/-^* compared with *bmf^+/+^* mice in the heart, lung, liver, or brain (data not shown). LC3-II accumulation was not observed in any tissue analyzed in *bmf^+/+^*, *bmf^+/-^*, and *bmf^-/-^* mice, even with higher doses of BAF (up to 50 µg per mice) at 6 or 24 hours post treatment (data not shown). These studies were conducted in 8 - 15 wks old mice. As back-up mechanisms may be more robust to mask effects of Bmf deficiency in young mice, we analyzed tissues from 56 wks old mice and found that LC3-II levels were decreased in the hearts of *bmf^+/-^* and even more in the heart of *bmf^-/-^* mice (**Figure 1G**). However, LC3-II levels in the lungs, kidneys, and spleens from 56 wks old mice still were not affected by Bmf deficiency (data not shown). Although LC3-II levels were decreased in the hearts of Bmf-deficient mice rather than increased, as observed in Bmf-deficient cells in culture, these observations support the overall idea that in aging conditions Bmf affects LC3-II degradation *in vivo*.

### Bmf inhibits autophagy independent of the mTOR pathway

In resting cells, the Ser/Thr protein kinase mTOR promotes protein translation (Ma & Blenis, 2009) while suppressing autophagy (Feng, Zhang et al., 2015). mTORC1 senses nutritional and environmental cues, and upon starvation is inhibited to activate autophagy (Nazio et al., 2013). To investigate whether the mTOR pathway is linked to the observed regulation of autophagy by Bmf, we inhibited mTOR by treating cells with pp242, a selective ATP-competitive inhibitor of mTOR, and analyzed the LC3 conversion in *bmf^+/+^*, *bmf^+/-^* and *bmf^-/-^* MEFs. mTOR activity was efficiently inhibited by pp242 in *bmf^+/+^* and *bmf^+/-^* MEFs, as detected by decreased amount of the phosphorylated form of P70S6K, a substrate for the mTORC1 kinase activity **(Figure S2A)**. This mTOR inhibition caused accumulated LC3-II in *bmf^+/+^*, *bmf^+/-^* and *bmf^-/-^* MEFs in the presence of BAF (**Figure S2A**). However, LC3-II levels in both vehicle- and pp242-treated cells were higher in *bmf^+/- (^***Figure S2A**, left blot) and in *bmf^-/-^* (**Figure S2A**, right blot) compared with *bmf^+/+^* MEFs and in *bmf^+/-^* compared with *bmf^+/+^* MAECs in the absence or presence of BAF (**Figure 2A**). Nutrient deprivation for 1 hour, another way to inhibit mTOR activity (Levine & Klionsky, 2004, Tasdemir, Maiuri et al., 2008), also effectively decreased the phospho-P70S6K levels, indicating mTOR inhibition (**Figure S2B**) and resulted in increased LC3-II accumulation in *bmf^-/-^* compared with *bmf^+/+^* MEFs in the presence of Baf (**Figure S2B**). The same results were observed in *bmf^+/-^* compared with *bmf^+/+^* MAECs after starvation (**Figure 2B**). Collectively, these results suggest that Bmf inhibits autophagy independent of the mTOR pathway.

**Figure 2:**
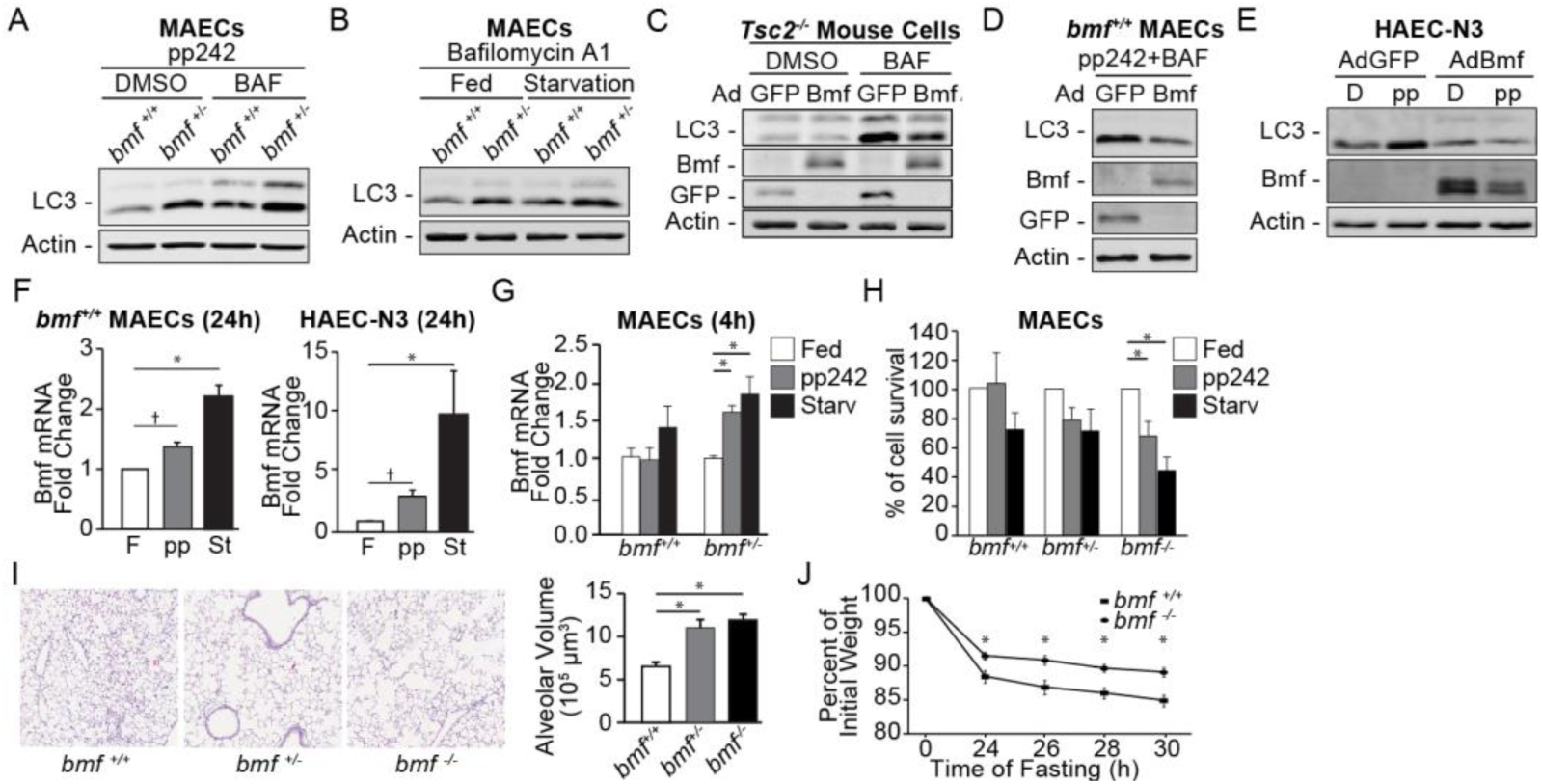
Bmf inhibits autophagy independent of the mTor pathway. **A)** Western blot analysis of protein extracts from *bmf^+/+^* and *bmf^+/-^* MAECs treated with pp242 (0.5 µM) and DMSO or BAF (100 nM) for 4 h. **B)** Western blot analysis of protein extracts from *bmf^+/+^* and *bmf^+/-^* MAECs in full media (Fed) or EBSS (Starvation) for 1 h in the presence of BAF (100 nM). **C)** Western blot of *Tsc2^-/-^* mouse cells infected with AdGFP or AdBmf (MOI: 50) for 24 h and treated with DMSO or BAF (50 nM) for the last 4 h. **D)** Western blot analysis of protein extracts from *bmf^+/+^* MAECs infected with AdGFP or AdBmf (MOI: 50) for 24 h and treated with pp242 (0.5 µM) for 4 h and BAF (100 nM) for the last hour. **E)** Western blot analysis of protein extracts from HAEC-N3 cells infected with AdGFP or AdBmf (MOI: 50) for 24 h and treated with DMSO or pp242 (2.5 µM) for the last 4 h. **F)** Quantification of Bmf mRNA from *bmf^+/+^* MAECs (left) or HAEC-N3 (right) treated with full media (F), pp242 (0.5 µM for MAECs and 2.5 µM for HAEC-N3) or EBSS (starvation, St) for 24 h and analyzed by qPCR (n= at least 3 independent experiments; ANOVA, * P<0.05). **G)** Quantification of Bmf mRNA from *bmf^+/+^* and *bmf^+/-^* MAECs treated with full media (Fed), pp242 (0.5 µM) or EBSS (starvation, Starv) for 4 h and analyzed by qPCR. (n= at least 3 independent experiments; ANOVA, * P<0.05). **H)** Quantification of viable *bmf^+/+^*, *bmf^+/-^* and *bmf^-/-^* MAECs after 24 h in full media (Fed), pp242 (0.5 µM) or EBSS (starvation, Starv) by Trypan blue exlusion assay (n= at least 3 independent experiments; ANOVA, * P<0.05). **I)** Representative images (left) and morphometric analysis for volume weighted mean alveolar volume (right) of lungs inflated and fixed with formalin under constant pressure, n= 3 mice per group (8 wks old male mice); ANOVA, * P<0.05. **J)** Percentage of body weight after fasting *bmf^+/+^* and *bmf^-/-^* mice for 30 h, relative to the body weight before start the fasting period, n= 6 mice per genotype (3 males and 3 females 9-10 wks old mice); unpaired T-test for each time point, * P<0.05.

Because Tsc2 protein is an endogenous mTOR inhibitor (Dibble & Cantley, 2015), *Tsc2^-/-^* cells have hyperactivation of mTOR as detected by high levels of phospho-P70S6K protein when compared with *Tsc2^+/+^* cells (**Figure S2C**). As expected, *Tsc2^-/-^* cells showed less LC3-II levels after BAF treatment compared with *Tsc2^+/+^* cells (**Figure S2C**). Therefore, we infected *Tsc2^-/-^* cells with AdGFP or AdBmf to investigate whether Bmf affects autophagy when mTOR is hyperactivated. We observed that BAF treatment of *Tsc2^-/-^* cells increased the LC3-II levels in control cells (AdGFP infected) and in cells expressing Bmf (AdBmf infected), but the accumulation of LC3-II was less when cells were expressing Bmf (**Figure 2C**), again supporting that Bmf inhibits autophagy irrespective of mTOR activation.

We further investigated whether Bmf overexpression reduces autophagy in different cell types when mTOR is inhibited. Therefore, MAECs were either infected with AdGFP or AdBmf and treated with pp242, to inhibit mTOR, and with BAF, to inhibit lysosomal degradation. Bmf over expression in *bmf^+/+^* MAECs reduced pp242-induced LC3-II accumulation in the presence of BAF (**Figure 2D**). Similar results were observed when Bmf levels were increased in *bmf^+/-^* or re-expressed in *bmf^-/-^* MAECs and cells were treated with pp242 and BAF (**Figure S2D**). When HAEC-N3 (**Figure 2E**) and HAEC-N1 (**Figure S2E**) were treated with pp242, the accumulated LC3-II levels were reduced in cells expressing Bmf compared with cells expressing GFP. These results suggest that Bmf inhibits autophagy downstream mTOR activation in murine and human cells.

Contrary to Bmf enhancing cell death, as was previously reported (Grespi, Soratroi et al., 2010, Labi et al., 2008, Puthalakath, Villunger et al., 2001), we had previously reported that survival of *bmf^-/-^* compared with *bmf^+/+^* MEFs is reduced when cells are starved or treated with pp242 for 24h (Contreras et al., 2013). Our observation suggested that by inhibiting autophagy, Bmf may protect from cell death induced by enhanced autophagy activation that occurs when mTOR is inhibited for prolonged periods. In support of this idea, Bmf mRNA expression was robustly increased both after pp242 treatment and starvation for 24h in wild type MAECs and in HAEC-N3 cells (**Figure 2F**). Further, starvation or pp242 treatment for 4 hours induced bmf mRNA levels by 1.5-2.0 fold in *bmf^+/-^* MAECs but not in *bmf^+/+^* MAECs, supporting the idea that increased Bmf expression may be needed to protect from cell death when mTOR is inhibited (**Figure 2G**). Additionally, bmf mRNA was also increased in *bmf^+/+^* MEFs by almost 4- and 8-fold after prolonged pp242 treatment and starvation, respectively (**Figure S2F**). Similar increases in Bmf mRNA were also observed in HAEC-N1 and AALEB cells after 24 hours of mTOR inhibition or starvation (**Figure S2G**). As expected, survival was reduced in *bmf^-/-^* compared with *bmf^+/-^* or *bmf^+/+^* MAECs (**Figure 2H**) in response to pp242 treatment or starvation over 24 hours. These findings further confirmed the protective role of Bmf expression from autophagy-induced cell death, likely by tempering the extent of autophagy induced by mTOR inhibition. Consistent with Bmf promoting survival, we observed extensive emphysema in the lungs of *bmf^-/-^* and *bmf^+/-^* compared with *bmf^+/+^* mice (**Figure 2I**). Therefore, persistent autophagy in the lungs may enhance death of alveolar cells in Bmf-deficient mice.

Our observations that Bmf may dampen autophagy when mTOR is inhibited suggested that fasting of Bmf-deficient mice may show a weight loss different than wild-type mice. B*mf^-/-^* mice showed reduced weight loss compared with *bmf^+/+^* mice after 30 h of fasting (**Figure 2J**).

### Bmf enhances proteasomal degradation

The observations that Bmf suppresses autophagy downstream mTOR lead us to investigate the underlying mechanisms for the autophagic inhibition by Bmf. It is known that a PI3 kinase needs to be activated downstream mTOR to trigger the initiation of autophagy. This PI3K activity requires Beclin1. Because we have previously observed that Beclin1 levels are increased in *bmf^-/-^* compared with *bmf^+/+^* MEFs and MAECs (Contreras et al., 2013), we wanted to test the hypothesis that Bmf deficiency may cause an imbalance between Beclin1 and Bcl-2 by increasing Beclin1 levels. Similar to what we previously reported in *bmf^-/-^* MAECs, Beclin1 levels were also higher in *bmf^+/-^* compared with *bmf^+/+^* MAECs; however, Bcl-2 protein levels were also higher in both *bmf^+/-^* and *bmf^-/-^* cells (**Figure 3A**). Therefore, we decided to investigate whether Bmf deficiency also increases the levels of other proteins. Interestingly, levels of proteins not related to autophagy regulation, including mitochondrial proteins like mitofusin 2 (Mfn2), cytochrome c (Cyt c) and Cox IV, the apoptotic protein Bak, the cytoskeleton protein dynein light chain 1/2 (DLC1/2), and the ER-associated protein BiP, were also higher in *bmf^-/-^* and *bmf^+/-^* compared with *bmf^+/+^* MAECs (**Figure 3A, and Figures S3A, B, C**). Another protein that was consistently increased in *bmf^+/-^* and *bmf^-/-^* MAECs was p62 (**Figures 3A** **and S3A)**. Beclin1, p62, and DLC1/2 were also elevated in HAEC-N3 and HAEC-N1 cells when Bmf levels were suppressed using shBmf (**Figure S3D**). The possibility that Bmf may affect protein stability was further strengthened by the observation that Beclin1 levels were reduced when Bmf levels were increased by AdBmf infection in *bmf^+/+^*, *bmf^+/-^* or *bmf^-/-^* MAECs (**Figure 3B**).

**Figure 3:**
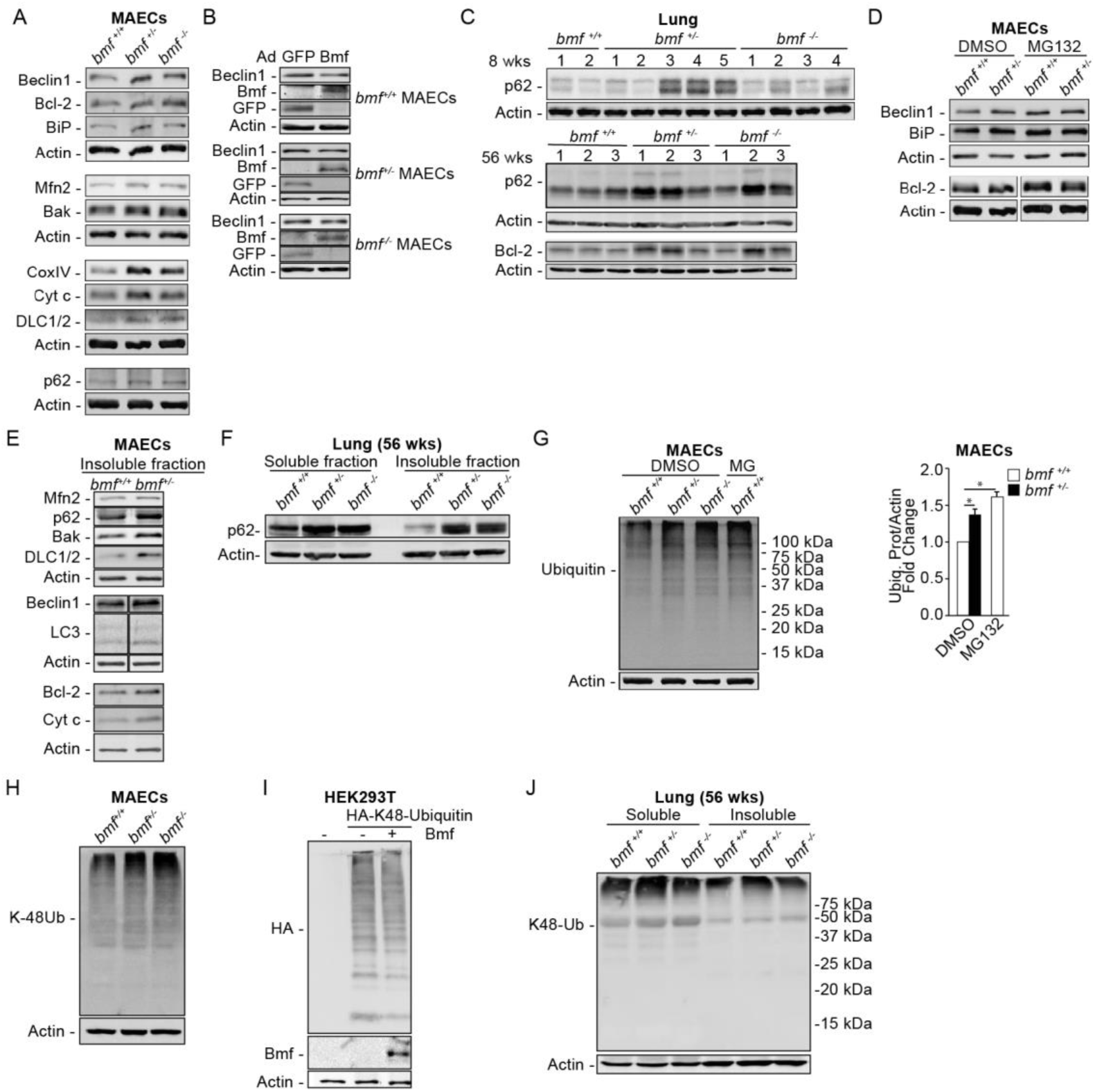
Bmf enhances proteasomal degradation. **A)** Western blot analysis of protein extracts from *bmf^+/+^*, *bmf^+/-^* and *bmf^-/-^* MAECs. **B)** Western blot analysis of protein extracts from *bmf^+/+^*, *bmf^+/-^* and *bmf^-/-^* MAECs infected with AdGFP or AdBmf for 24 h (MOI: 50). **C)** Western blots analysis of protein extracts from the lung cranial lobe lysates (100 µg of proteins) from male mice at 8 and 56 weeks of age. **D)** Western blots analysis of protein extracts from *bmf^+/+^* and *bmf^+/-^* MAECs treated with DMSO or MG132 (10 µM) for 6 h. **E)** Western blots of the “insoluble” fraction from *bmf^+/+^* and *bmf^+/-^* MAECs. **F)** Western blot of the “soluble” and “insoluble” fractions from the left lung (100 µg of proteins) from 56 wks old male mice. **G)** Western blot of *bmf^+/+^*, *bmf^+/-^* and *bmf^-/-^* MAECs treated with DMSO and *bmf^+/+^* MAECs treated with MG132 (10 µM) for 6 h (left) and graph showing ubiquitinated proteins/Actin fold change, n= at least 3 independent experiments (right); ANOVA, * P<0.05. **H)** Western blot of *bmf^+/+^*, *bmf^+/-^* and *bmf^-/-^* MAECs. **I)** Western blot of HEK293T cells either not transfected, transfected with HA-K48-Ubiquitin, or with HA-K48-Ubiquitin and Bmf_S_-expressing plasmids. **J)** Western blot of the “soluble” and “insoluble” fractions from the left lung from 56 wks old male mice and probed for K48-Ubiquitin and actin.

As was observed in *bmf^+/-^* and *bmf^-/-^* MAECs (**Figure 3A**), p62 levels were also increased in the lungs of *bmf^+/-^* and *bmf^-/-^* mice at 8 wks (**Figure 3C**) and this p62 increase was more evident in *bmf^-/-^* mice at 15 wks of age (**Figure S3E**). BAF did not modify p62 levels in the tissues analyzed (data not shown), but age robustly increased p62 levels (**Figure S3F**) and the lungs, hearts, kidneys and spleens from *bmf^+/-^ or bmf^-/-^ s*how higher levels of p62 compared with *bmf^+/+^* mice when mice are 56 wks old (**Figures 3C, S3G**). Bmf deficiency caused a significant increase of p62 *in vivo* than in cultured cells (**Figures 3A and 3C**). Increased p62 levels were also evident in the heart, liver and muscle from *bmf^-/-^* mice compared with *bmf^+/+^* mice that were fasted for 30h (**Figure S3H**). We found that other proteins were also elevated in different tissues from *bmf^+/-^* mice at 8 or at 15 weeks (data not shown). Most of the proteins analyzed (including Bcl-2, CoxIV and BiP) were detected to be at even higher levels in Bmf-deficient lungs, hearts, kidneys and spleens from 56 weeks old mice (**Figures 3C, and S3I**).

As Bmf having a role in protein stability has not been reported before, we decided to explore this role by treating *bmf^+/-^* and *bmf^+/+^* MAECs with MG132. We observed accumulation of several proteins in *bmf^+/+^* but not in *bmf^+/-^* MAECs, consistent with an inhibition of their proteasomal degradation (**Figures 3D, and S3J, S3K).** These observations suggested that proteasomal degradation is disrupted in Bmf-deficient cells. When proteasomal degradation is inhibited, the accumulated proteins are confined in inclusion bodies as aggregates to decrease their cell toxicity (Hillert, Brnjic et al., 2019, Takahashi, Kitaura et al., 2018). Therefore, we investigated whether *bmf^+/-^* MAECs present with accumulation of proteins in the “insoluble fraction” that is solubilized only with high concentrations of SDS and DDT. We found that *bmf^+/-^* compared with *bmf^+/+^* MAECs show increased accumulation of several proteins also in the “insoluble fraction” (**Figures 3E** **and S3L**). We also found that the “insoluble fractions” from the Bmf-deficient lungs (**Figure 3F**), liver, and muscle (data not shown) present with increased p62 compared with wild-type mice. Also, *bmf^+/+^* MAECs present with more proteins in the “insoluble fraction” after treatment with MG132 (**Figure S3M**), confirming that it is a phenotype associated with proteasomal degradation inhibition. These observations suggest that Bmf facilitates proteasomal degradation.

In another set of experiments, we compared levels of total poly-ubiquitinated proteins in *bmf^+/+^*, *bmf^+/-^*, and *bmf^-/-^* MAECs. Poly-ubiquitinated proteins were increased in *bmf^+/-^* and *bmf^-/-^* compared with *bmf^+/+^* MAECs (**Figure 3G**). Also, wild type MAECs treated with MG132 accumulated poly-ubiquitinated proteins to levels observed in non-treated *bmf^+/-^* MAECs (**Figure 3G**), further supporting the idea that Bmf enhances proteasomal degradation. We also found that *bmf^+/-^* and *bmf^-/-^* compared with *bmf^+/+^* MAECs present with more K-48 poly-ubiquitination (**Figure 3H**), the prominent ubiquitination associated with proteasomal degradation.(Yau & Rape, 2016) When HEK293T cells were co-transfected with HA-K48-ubiquitinin and Bmf, the total K48-poly-ubiquitination of proteins was reduced compared with cells not transfected with Bmf as control (**Figure 3I**). Also, the lungs from *bmf^+/-^* or *bmf^-/-^* compared with *bmf^+/+^* mice showed more total K-48-polyubiquitinated proteins both in the “soluble” and “insoluble” fractions (**Figure 3J**). Together, these findings point to the role of Bmf facilitating proteasomal degradation *in vitro* and *in vivo*.

### Bmf affects degradation of mucins in mice and in differentiated MAECs

We decided to investigate whether the role of Bmf in inhibiting protein degradation *in vivo* will be more evident following an injury to the lung. We have previously shown that in mice, prolonged exposure to ovalbumin (OVA) as an allergen causes the resolution of mucous cell hyperplasia (Shi, Fischer et al., 2002, Tesfaigzi, Fischer et al., 2002). Consistent with previous observations, the number of mucous cells was reduced in *bmf^+/+^* mice 15 days after the OVA exposure, but resolution was abrogated in *bmf^-/-^* mice (**Figure 4A**). Because of our findings that Bmf enhances proteasomal degradation, we decided to investigate whether Bmf plays a role in reducing stored muco-substances by proteolytic degradation of mucins. Using differentiated MAECs, we first established an *in vitro* system that reflects what was observed in mice. Therefore, *bmf^+/+^* and *bmf^+/-^* MAECs were differentiated on air-liquid interface cultures and treated with IL-13 for 14 d to induce mucin production. Another set of cultures were treated with IL-13 for 14 days and kept without treatment for an additional 7 days. As detected by immunofluorescence, *bmf^+/-^* compared with *bmf^+/+^* MAECs showed more Muc5ac-positive cells after 7 days of recovery post IL-13 treatment (**Figure 4B**). In addition, although Muc5ac mRNA levels were similar in untreated *bmf^+/+^*, *bmf^+/-^* and *bmf^-/-^* differentiated MAECs (**Figure 4C**), Western blot analysis of the mucins first separated in a 0.8% agarose gel, transferred to a nitrocellulose membrane and developed with antibodies anti Muc5ac showed that *bmf^-/-^* compared with *bmf^+/+^* MAECs displayed higher Muc5ac protein levels both at baseline and at 7 d post IL-13 treatment (**Figure 4D**). In addition, mucins in *bmf^-/-^* compared with *bmf^+/+^* MAECs showed a higher signal with anti-ubiquitin antibody, (**Figure 4D**). Also, antibodies to K48-ubiquitin showed a higher signal in mucins from *bmf^-/-^* compared with *bmf^+/+^* MAECs under control or IL-13 stimulated conditions (**Figure 4E**), suggesting that mucin degradation may be delayed by Bmf deficiency. Similarly, differentiated *bmf^+/-^* MAECs showed more Muc5ac and more K48-ubiquitinated mucins than wild type cells at baseline (**Figure S4A**) and after IL-13 treatment (**Figure S4B**). Differentiated *bmf^-/-^* compared with *bmf^+/+^* MAECs also showed accumulation of other proteins (**Figure 4F**). As expected, differentiated *bmf^+/-^* MAECs showed accumulation of several proteins compared with *bmf^+/+^* MAECs at baseline (**Figure S4C**) and after IL-13 treatment (**Figure S4D**). IL-13 treatment reduced Bmf mRNA levels in both differentiated *bmf^+/+^* and *bmf^+/-^* MAECs (**Figure S4E**), supporting the idea that reduced Bmf is correlated with increased Muc5ac protein levels.

**Figure 4:**
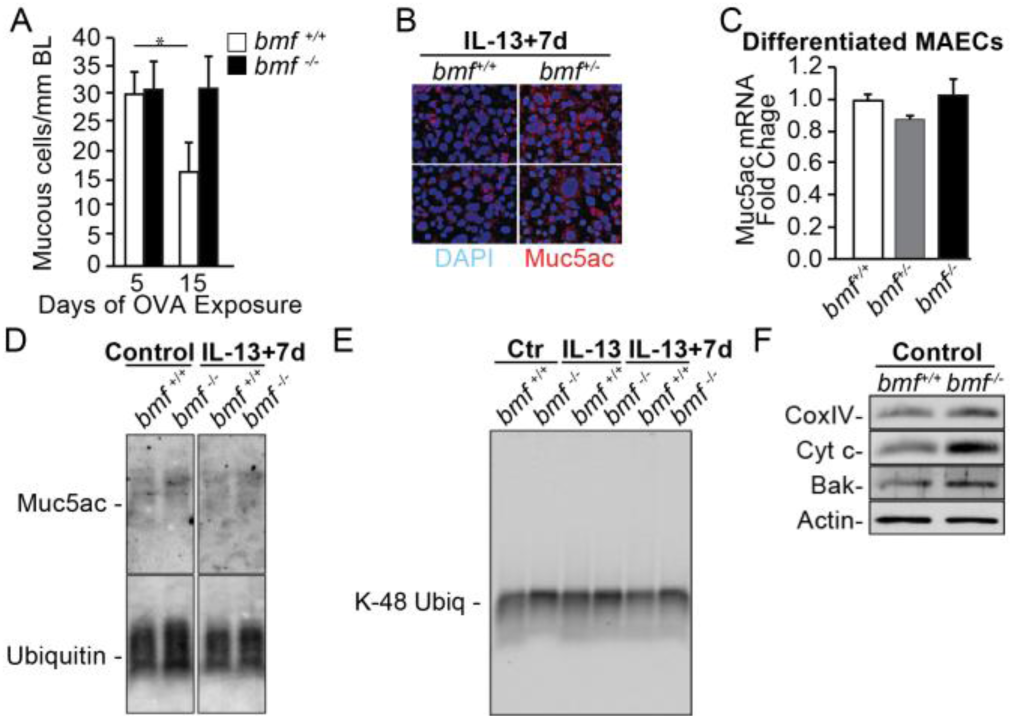
Bmf affects proteasomal degradation of mucins in mice and in differentiated MAECs. **A)** Mucous cells per mm of basal lamina in lungs from *bmf^+/+^* and *bmf^-/-^* mice after 5 or 15 days of OVA exposure, n= 5 mice per genotype (4 males and 1 female *bmf^+/+^* and 2 males and 3 females *bmf^-/-^* mice); unpaired T-test, * P<0.05. **B)** Immunofluorescence for Muc5ac in differentiated *bmf^+/+^* and *bmf^+/-^* MAECs treated with IL-13 (10 ng/ml) for 14 days and kept without treatment for an additional 7 days period. **C)** Quantification of Muc5ac mRNA from differentiated *bmf^+/+^*, *bmf^+/-^* and *bmf^-/-^* MAECs in control media and analyzed by qPCR. **D)** Protein extracts prepared from differentiated *bmf^+/+^* and *bmf^-/-^* MAECs in control media (control) or 7 days in control media post 14 days of 10ng/ml IL-13 treatment (IL-13 + 7d) and analyzed by Western blotting after agarose gel electrophoresis using anti Muc5ac and anti Ubiquitin antibodies. **E)** Protein extracts prepared from differentiated *bmf^+/+^* and *bmf^-/-^* MAECs in control media (control), treated with 10ng/ml IL-13 for 14 days (IL-13) or 7 days post IL-13 treatment (IL-13 + 7d) and analyzed by Western blotting after agarose gel electrophoresis using anti K-48-Ubiquitin antibodies. **F)** Western blot analysis of protein extracts from differentiated *bmf^+/+^* and *bmf^-/-^* MAECs in control media and developed with anti CoxIV, Cytochrome c (Cyt c), Bak and Actin antibodies.

### Bmf inhibits autophagy by decreasing Beclin-K63 ubiquitination

Because our findings strongly suggest that Bmf facilitates proteasomal degradation both *in vitro* and *in vivo*, we explored the hypothesis that Bmf may be linking the proteasomal degradation and the autophagy pathways. We investigated whether Bmf inhibits autophagy by stabilizing the Beclin1/Bcl-2 complex by preventing Beclin1 from being released in its activate form to initiate the autophagosome formation (Pattingre et al., 2005). Immunoprecipitation with anti-Beclin1 antibodies resulted in reduced Bcl-2 protein from *bmf^+/-^* (heterozygote) compared with *bmf^+/+^* MAEC extracts, confirming that Bmf stabilizes the Beclin1/Bcl-2 complex (**Figure 5A**). Similarly, reduced Bcl-2 binds Beclin1 in *bmf^-/-^* (knockout) compared with *bmf^+/+^* MAECs (**Figure S5A**), suggesting that Beclin1 is released from Bcl-2 as Bmf levels are reduced.

**Figure 5:**
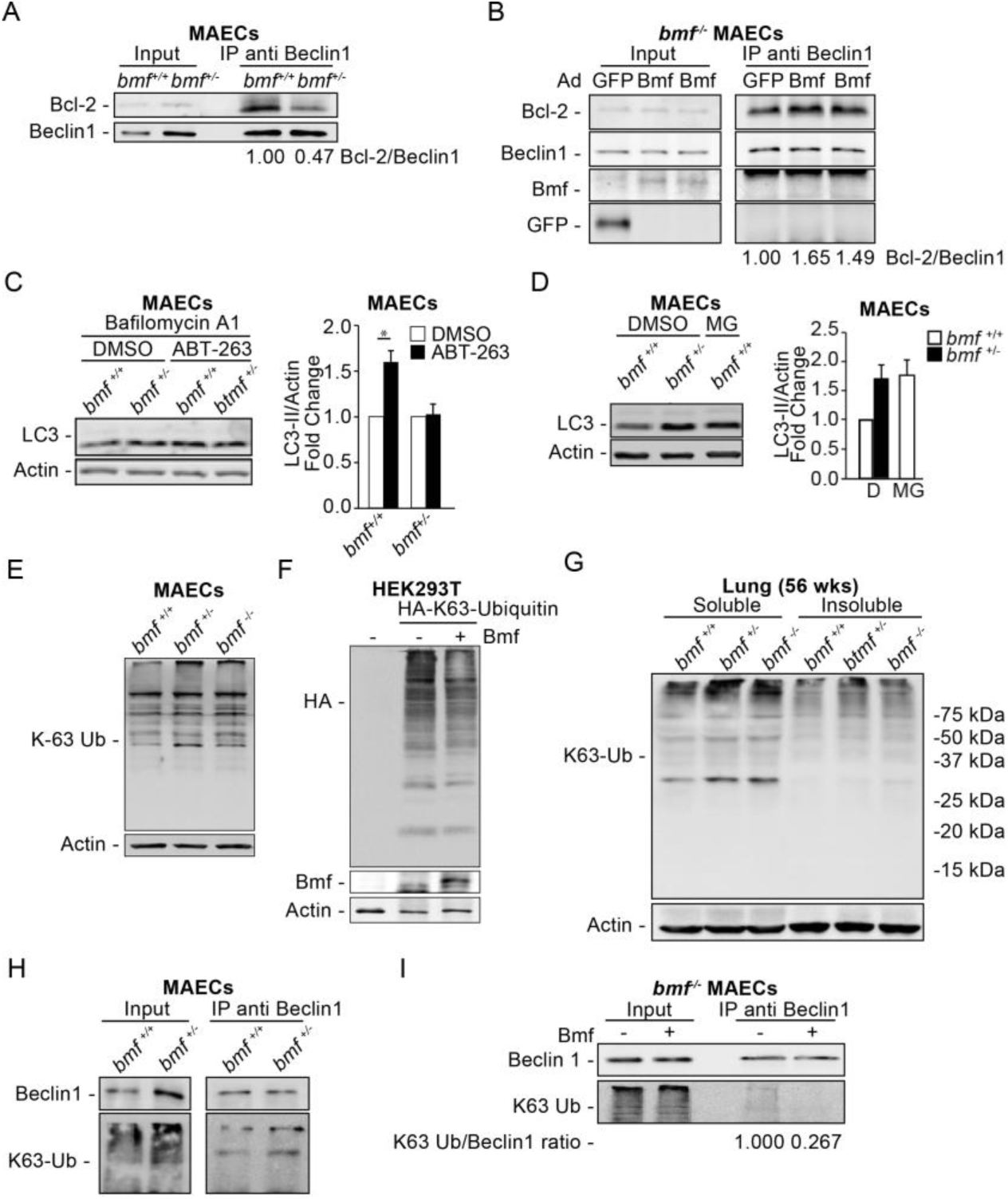
Bmf inhibits autophagy by decreasing K63 ubiquitination of Beclin1. **A)** Immunoprecipitation of protein extracts from *bmf^+/+^* and *bmf^+/-^* MAECs using anti Beclin1 antibodies and Western blot analysis of inputs and immunoprecipitated (IP) proteins. The ratio Bcl-2/Beclin1 in the immunoprecipitates is shown. **B)** Immunoprecipitation of protein extracts from *bmf^-/-^* MAECs infected with AdGFP or AdBmf (MOI: 100) for 24 h using anti Beclin1 antibodies. Western blot analysis of inputs and immunoprecipitated (IP) proteins with calculated Bcl-2/Beclin1 ratio. **C)** Western blot of *bmf^+/+^* and *bmf^+/-^* MAECs treated with ABT-263 (0.5 µM) for 5 h and BAF (100 nM) for the last hour (left) and graph showing the fold change of LC3-II/Actin, n= at least 3 independent experiments (right); ANOVA, * P<0.05. **D)** Western blot analysis of protein extracts from *bmf^+/+^* and *bmf^+/-^* MAECs treated with DMSO or MG132 (10 µM) for 6 h (left) and graph showing the fold change of LC3-II/Actin, n= at least 3 independent experiments (right). **E)** Western blot analysis of *bmf^+/+^*, *bmf^+/-^* and *bmf^-/-^* MAECs for total K-63 ubiquitinated proteins and actin. **F)** Proteins extracted from non-transfected HEK293T cells and those transfected with HA-K63-Ubiquitin or co-transfected with both HA-K63-Ubiquitin and Bmf_S_ expressing plasmids and analyzed by Western blotting. **G)** Western blot analysis of the “soluble” and “insoluble” fractions from the left lung of male mice at 56 wks of age. **H)** Immunoprecipitation of protein extracts from *bmf^+/+^* and *bmf^+/-^* MAECs using anti Beclin1 antibodies. Western blot analysis of inputs and immunoprecipitated (IP) proteins. **I)** Immunoprecipitation using anti Beclin1 antibodies of protein extracts from *bmf^-/-^* MAECs non transfected or transfected with a Bmf_S_ expressing plasmid, inputs and immunoprecipitated (IP) proteins analyzed by Western blot. The calculated K63-Ubiquitin/Beclin1 ratio is shown.

We have previously detected the interaction of Bmf with Beclin1 and Bcl-2 in wild type MEFs (Contreras et al., 2013). As endogenous Bmf is only faintly detected by Western blotting in extracts prepared from MAECs (**Figure S1A**), we decided to increase the expression of Bmf using adenoviral expression vector in *bmf^-/-^* MAECs and analyze Beclin1-interacting proteins by immunoprecipitation compared with Ad-GFP-infected controls. The Beclin1/Bcl-2 interaction was more abundant in cells expressing Bmf (**Figure 5B**). However, although Bmf was expressed, as shown in the input, Bmf protein was not detectable in the immunoprecipitates with anti-Beclin1 antibodies. Similarly, infecting HEK293T cells with AdBmf to increase Bmf expression resulted in increased interaction of Beclin1/Bcl-2 when the IP is performed with anti-Bcl-2 antibodies (data not shown), although Bmf was not detected in the IP. These findings may suggest that Bmf easily dissociates from the complex during the immunoprecipitation process.

Because inhibition of autophagy occurs when Beclin1 interacts with the endoplasmic reticulum (ER)-localized Bcl-2 (Pattingre et al., 2005), we designed an experiment overexpressing ER-localized Bcl-2 (ER Bcl-2) by transiently transfecting HEK293T cells and 2 h later infecting them with AdBmf. Immunoprecipitation with anti-Bcl-2 antibodies showed that Bmf enhances ER-Bcl-2 interaction with Beclin1 (**Figure S5B)**, and Bmf was detected in the IP revealing the interaction of Bmf with ER-localized Bcl-2.

We further investigated whether autophagy is activated in Bmf-deficient cells by promoting Beclin1 release from Bcl-2. We used the BH3 mimetic ABT-263 that releases Beclin1 from Bcl-2 due to its higher affinity to Bcl-2.(Maiuri, Zalckvar et al., 2007) ABT-263 treatment increased LC3-II levels in *bmf^+/+^* but not in *bmf^+/-^* MAECs (**Figure 5C**), supporting the idea that activation of autophagy may occur by releasing Beclin1 from Bcl-2 either by the presence of a BH3 mimetic compound or by reduced Bmf levels. The involvement of the proteasomal activity by Bmf on autophagy regulation was examined by using MG132. Upon 6h of MG132 treatment, LC3-II increased in *bmf^+/+^* MAECs to levels similar to what was observed in *bmf^+/-^* MAECs treated with DMSO as vehicle control (**Figure 5D**). As MG132 blocks proteasomal function, these findings further support the idea that the Beclin1/Bcl-2 interaction was affected by Bmf-deficiency due to the inhibition of the proteasome.

When the proteasomal degradation is disrupted, certain E3 ligases increase the K63-linkage ubiquitination of many proteins.(Lim, Chew et al., 2013) When Beclin1 is ubiquitinated by K63-linkage, the Beclin1/Bcl-2 interaction is disrupted and Beclin1 is released to activate the PI3P kinase activity and autophagy is activated.(Shi & Kehrl, 2010a) Therefore, we investigated whether this process may be responsible for the activated autophagy in Bmf-deficient cells. Overall, K63-linked ubiquitination is increased in *bmf^-/-^* and *bmf^+/-^* compared with *bmf^+/+^* MAECs (**Figure 5E**). K63-linked ubiquitination is also increased in *bmf^-/-^* compared with wild type MEFs (**Figure S5C**). Also, HEK293T cells that co-express HA-K63-Ubiquitin and Bmf show reduced K63-poly-ubiquitination compared with cells not overexpressing Bmf (**Figure 5F**). In addition, K-63-polyubiquitinated proteins are increased in the lungs of *bmf^+/-^* or *bmf^-/-^* compared with *bmf^+/+^* mice in both the “soluble” and “insoluble fractions” (**Figure 5G**).

Immunoprecipitation with anti-Beclin1 antibodies confirmed increased K63-linked ubiquitination of Beclin1 in *bmf^+/-^* compared with *bmf^+/+^* MAECs (**Figure 5H**). Further, immunoprecipitation of Beclin1 from *bmf^-/-^* MAECs transfected with a Bmf-expression plasmid showed reduced K63-ubiquitination of Beclin1 (**Figure 5I**). However, likely due to the low transfection efficiency in MAECs, expressed Bmf was not detected by Western blotting. Collectively, these results demonstrate that in MAECs, Bmf-deficiency not only increases Beclin1 levels but also facilitates K63 ubiquitination of Beclin1 to cause the release of Beclin1 from Bcl-2 and thereby activate autophagy.

### Dynein Binding and BH3 Domains of Bmf affect K63-ubiquitination and autophagy

To identify the Bmf protein domains responsible for reducing K63-ubiquitination, Bmf mutants in the dynein-binding domain (DBD mutant) or the BH3 domain (BH3 mutant) were generated within the Bmf_S_ isoform (**Figure 6A**). Transfected HEK293T cells showed two Bmf bands, with the BH3 mutant expressing more of the larger band and both the DBD mutant and the wild type expressing more of the smaller band (**Figure 6B**). When co-transfected along with HA-K63-ubiquitin expression plasmid, both the DBD and BH3 mutants were less efficient in reducing total K63-ubiquitinaton than wild-type Bmf (**Figure 6C**). Immunoprecipitations with anti Beclin1 antibodies showed reduced K63-ubiquitinated Beclin1 in cells transfected with wild-type compared with Bmf mutants (**Figure 6D**). Also, the Beclin1/Bcl-2 interaction increased more by wild-type Bmf than by the Bmf mutants (**Figure 6D**). Because we did not observe changes in LC3-II between HEK293T cells expressing wild-type or mutant Bmf forms, we suspected that endogenous Bmf in HEK293T cells may play a role. Therefore, we transfected *bmf^-/-^* MEFs with these constructs and detected Bmf only in the “insoluble fraction”; also, Bmf mRNA was detected only in cells transfected with w.t., BH3, and DBD mutants (**Figure S6A**). Expression of Bmf was confirmed by detecting Bmf mRNAs by qPCR (data not shown). Despite low Bmf protein levels detected, cells transfected with wild-type Bmf showed decreased BAF-induced LC3-II accumulation compared with cells transfected with the BH3 or DBD mutants, with the DBD mutant showing the highest LC3-II levels (**Figure 6E**). We have shown previously that *bmf^-/-^* MEFs present higher numbers of LC3 puncta than wild type MEFs when overexpressing LC3-Cherry (Contreras et al., 2013). While *bmf^-/-^* MEFs also present with a high number of LC3 puncta (**Figure 6F**), the number of LC3 puncta (**Figure 6F**) as well as their intensity (**Figure S6B**) were reduced when cells were transfected with wild-type Bmf but not with the BH3 or DBD mutant forms. These results suggest that the BH3 and DBD domains of Bmf are required for the effects of Bmf on protein ubiquitination and autophagy.

**Figure 6:**
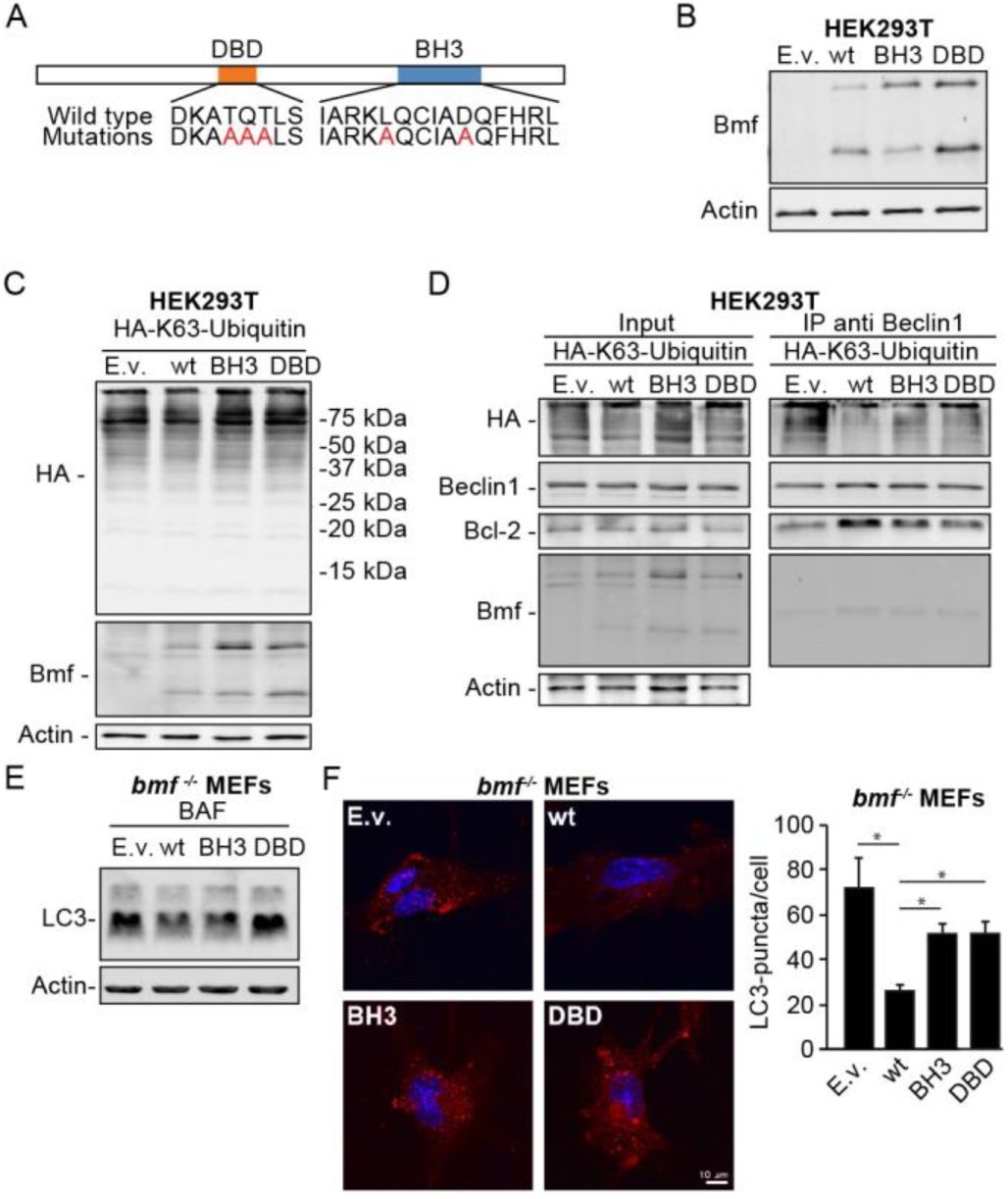
Domains of Bmf protein important for its role on K63-ubiquitination and autophagy. **A)** Amino acid sequences of the dynein binding domain (DBD) and the BH3 only domain of the mouse wild type Bmf_S_ protein and the introduced amino acids mutations in the DBD and the BH3 mutants. **B)** Western blot analysis of protein extracts from HEK293T cells transfected with control plasmid (Empty vector, E.v.), or those expressing wild type Bmf_S_ (wt), and the BH3 or DBD mutants. **C)** Proteins from HEK293T cells co-transfected with HA-K63-Ubiquitin and control (Empty vector, E.v.) or Bmf_S_ (wild type and mutants) expressing plasmids were analyzed by Western blotting. **D)** Immunoprecipitation of protein extracts from HEK293T cells co-transfected with HA-K63-Ubiquitin and E.v., wild type (wt) Bmf_S_ or BH3 and DBD mutants using anti Beclin1 antibodies. Western blot analysis of inputs and immunoprecipitated (IP) proteins. **E)** *Bmf^-/-^* MEFs transfected with E.v., wt, or the BH3 and DBD mutants plasmids developed with anti LC3 and anti Actin antibodies. F) Representative immunofluorescent images of *bmf^-/-^* MEFs transfected with the E.v, wt, or BH3 and DBD mutant plasmids and immunostained with anti LC3 and Cy5 antibodies (left) and quantification of punctuated LC3 of 23-48 cells from each transfection (ANOVA, * P<0.05).

## DISCUSSION

The present study shows that the main role of Bmf is to facilitate the degradation of many proteins and Bmf-deficiency, by increasing levels of ubiquitinated proteins causes reduced protein degradation and increased autophagy. Therefore, in Bmf deficient cells, ubiquitination of Beclin1 by the K63-linked chain causes the release of Beclin1 from Bcl-2 to initiate autophagy. This finding explains the seemingly contradictory role of Bmf as a BH3-only protein, rather than competing for the BH3 binding groove of Bcl-2, enhances the Bcl-2/Beclin1 interaction.

Starvation of cells causes Beclin1 to be ubiquitinated at Lys437 via a K63 linkage. Beclin1 is also K63-ubiquitinated after TLR4 stimulation in macrophages, but this ubiquitination occurs on Lys117.(Shi & Kehrl, 2010b) In contrast, WASH negatively regulates autophagy through suppression of Beclin1 K63-polyubiquitination.(Xia, Wang et al., 2013). Phosphorylation of USP14 by Akt removes K63 ubiquitin linkages including that of Beclin1.(Xu, Shan et al., 2016) Whether Bmf participates in deubiqutinating both sites on Beclin1 will be studied in the future.

Mutations on the BH3 or DBD domains of Bmf compromise the K63 ubiquitination of Beclin1. It is possible that the BH3 and DBD mutations affect Bmf function because of conformational changes. However, because the DBD attaches Bmf to the actin-myosin V filaments through DLC2, it is possible that Bmf may act as a sensor for stimulators of the cytoskeletal structures to regulate protein degradation. Several deubiquitinases (DUBs) have been associated with the cytoskeletal machinery of the cell, including USP33 (Li, D’Angiolella et al., 2013) and ataxa-3.(Burnett & Pittman, 2005)

This report focused on mutations of Bmf_S_, but did not explore the Bmf_L_ or Bmf_CUG_ isoforms. Whether these protein variants, that originate from different transcriptional start sites. may determine the cell death or proteasomal roles of Bmf needs further investigation. It is possible that the relative level of each Bmf isoform determines its role in protein degradation and autophagy, generating an effect that is tissue- and cell type-dependent.

Interestingly, *bmf^+/-^* compared with *bmf^-/-^* MAECs present with more Beclin1 levels; therefore, most of our studies used *bmf^+/-^* cells. Also, the inhibitory role of Bmf on autophagy becomes evident only under stress conditions, such as cells maintained in dispersed culture conditions. Surprisingly, when MAECs were differentiated on Transwell cultures the levels of several protein were increased in Bmf deficient cells, but Beclin1 levels remained unchanged in *bmf^-/-^* and *bmf^+/-^* compared with *bmf^+/+^* genotypes. Similarly, the levels of Beclin1 in the lungs or other tissues were not different among *bmf^+/+^* and *bmf^+/-^* or *bmf^-/-^* mice (data not shown). Autophagy, as detected by changes in the LC3-II levels, was not observed in most tissues of Bmf-deficient mice, except for in the heart of old mice, where LC3-II levels were reduced in the Bmf-deficient mice. These findings suggest that compensatory mechanisms that reduce Beclin1 ubiquitination may be activated when Bmf is absent. Fasting in mice does not induce autophagy uniformly in all tissues, suggesting that regulation of autophagy is organ specific.(Yoshimoto, Shibata et al., 2014) Accordingly, while accumulation of p62 in Bmf-deficient mice was detected in all tissues analyzed, autophagy phenotype by Bmf deficiency was detected only in the heart.

The fact that Bmf deficiency increases LC3-II levels in cultured cells but reduces LC3-II levels in heart tissue may be explained by an enhanced autophagic flux in mouse tissues. Also, we found reduced weight loss during fasting of *bmf^-/-^* compared with *bmf^+/+^* wild-type mice. In contrast, previous findings show that reduced autophagy is associated with reduced weight loss in fasting mice.(Fernandez, Barcena et al., 2017) Therefore, reduced weight loss during fasting in Bmf deficient mice may be a result of the disrupted protein degradation, the phenotype that is predominantly observed in many tissues. Future studies will investigate the role of protein degradation and weight loss during fasting.

The observation that Bcl-2, DLC1/2, BiP, Mfn2 and p62 levels are increased in several tissues (spleen, brain, thymus, lung) from *bmf^+/-^* compared with *bmf^+/+^* mice as young as 8-15 wks old suggests that the major function of Bmf *in vivo* is to regulate protein degradation. The unfolded protein response (UPR) is crucial for selectively degrading misfolded proteins.(Grice & Nathan, 2016) When the UPR is overloaded to degrade the K48-tagged proteins, K63–linked ubiquitination initiated as a signal to promote sequestration of proteins into aggresome.(Olzmann & Chin, 2008) Given that K63-linked polyubiquitination of proteins is generally uncoupled from the UPR, enhanced cellular ubiquitin modification of proteins via K63 promotes their accumulation and subsequent aggregation in the cell (Lim et al., 2013). We also found that Bmf deficiency enhances aggresome formation, further supporting that Bmf reduces K63-linked polyubiquitination.

The most striking phenotype of the Bmf-deficient mice is the increase in p62, as it was observed in cultured cells, but even more prominently in all tissues analyzed. Accumulation of p62 is delayed in *bmf^-/-^* compared with *bmf^+/-^* mice and fasting did not affect the higher levels of p62 in the heart, muscle and liver of *bmf^-/-^* compared with *bmf^+/+^* mice, both in the “soluble” and “insoluble” fractions. Proteasome inhibition induces rapid p62 expression in cells, which enhances survival by sequestering ubiquitinated proteins in inclusion (Sha, Schnell et al., 2018) and to sequester and attach the cargo to LC3-tagged isolation membranes for phagosome formation (Zaffagnini, Savova et al., 2018). Accumulation of p62 in Bmf-deficient mice remains unaffected when fasting over 30h, suggesting that the role of Bmf in proteasomal degradation inhibition is not modified by inhibiting the mTORC1 pathway. The observation that generally ubiquitinated proteins are increased by Bmf-deficiency suggests that the protein degradation system is affected by Bmf deficiency to such an extent that in addition to increased autophagy sequestration of proteins in aggresomes may be needed by increasing p62 protein levels. How Bmf deficiency leads to increased p62 levels is not known.

Metaplastic mucous cells and the amount of stored mucosubstances remained unchanged in *bmf^-/-^* mice and did not resolve following 15 d of challenge with ovalbumin, while the number of metaplastic mucous cells were reduced in *bmf^+/+^* mice. Our findings suggest that the metaplastic mucous cells remained because of lack of degradation of mucins. We found that more ubiquitinated mucins are present in *bmf^-/-^* and *bmf^+/-^* compared with *bmf^+/+^* differentiated MAEC cultures, suggesting that mucins may undergo degradation even under normal conditions. Intracellular levels of mucins are believed to be regulated by secretion. (Innes, Woodruff et al., 2006, Roy, Livraghi-Butrico et al., 2014, Singer, Martin et al., 2004) Also, in airway epithelial cells, IL13 activates autophagy to regulate secretion.(Dickinson, Alevy et al., 2016) This is the first report to show that mucins are ubiquitinated. Mucins are stored in vesicles tethered to myosin fibers (Lin, Fang et al., 2010, Raiford, Park et al., 2011) and we showed that the ubiquitination of mucins is increased when Bmf is deficient. Because mucins are highly glycosylated proteins, ubiquitination and degradation of these complex molecules requires E ligases that transfer the ubiquitin to these huge glycoproteins. An E3 ubiquitin ligase that recognizes sugar chains, Fbx2, that is primarily found in neuronal cells (Yoshida, Chiba et al., 2002) and one that recognizes *N*-glycan, Fbs2 (called Fbx6b or FBG2 previously) (Yoshida, Tokunaga et al., 2003) have been identified. Approximately 70 genes for F-box proteins, and at least five homologous F-box proteins containing a conserved motif in their C-termini are thought to recognize sugar chain of N-linked glycoproteins.(Yoshida & Tuder, 2007) Among these, Fbs1 and Fbs2 are perhaps involved in the endoplasmic reticulum-associated degradation pathway. Whether these proteins are involved in the degradation of mucins remains to be investigated. However, modifying the degradation of mucins may provide a novel approach to reduce epithelial stored mucosubstances in chronic lung diseases before their secretion obstructs the airways.

We found that Bmf-deficient mice present with emphysema already at 8 wks of age. Cigarette smoke, by inducing PINK, mediates autophagy-dependent elimination of mitochondria (mitophagy) and necroptosis of alveolar epithelial cells to cause emphysema in mice (Mizumura, Cloonan et al., 2014). Another study found that cigarette smoke causes cell death of alveolar cells by decreasing Bcl-2/Beclin1 interaction and inducing autophagy.(Qin, Gao et al., 2019) In fact, basal level of autophagy is always present in type II alveolar cells and is increased after a fasting period (Mizushima, Yamamoto et al., 2004). Transgenic expression of IFNγ in airways driven by the CCSP promoter causes emphysema (Wang, Zheng et al., 2000) and we showed that IFNγ causes autophagy in airway epithelial cells by suppressing Bmf expression.(Contreras et al., 2013) Human alveolar epithelial cells treated with cigarette smoke extract have increased polyubiquitinated proteins and cell death, and mice exposed to cigarette smoke show impaired proteasomal activity and accumulation of polyubiquitinated proteins in the soluble and insoluble protein fractions.(van Rijt, Keller et al., 2012) Further, p62 levels are higher in smokers with severity of emphysema.(Tran, Ji et al., 2015) Together with the findings in the current study that reduced Bmf levels increase p62 levels and enhance the autophagic cell death of type II cells, we postulate that increasing Bmf expression in lung epithelial cells may be an effective approach to block the development of emphysema.

## Materials and Methods

### Animals

Pathogen free *bmf^-/-^* mice on the C57BL/6 background were previously described (Labi et al., 2008) and were made available by A. Strasser (The Walter and Eliza Hall Institute of Medical Research, Melbourne, Australia). These mice along with the wild-type were bred at the Lovelace Respiratory Research Institute (LRRI) under specific pathogen-free conditions and genotyped as described previously (Labi et al., 2008). All animal experiments were approved by the Institutional Animal Care and use Committee (IACUC) and were performed at LRRI, a facility approved by the Association for the Assessment and Accreditation for Laboratory Animal Care International. Mice were injected intraperitoneally with DMSO (Sigma) or bafilomycin A1 (Santa Cruz) at doses of 0.1 or 2.5 mg/Kg, and 6 or 24 h later they were humanely euthanized for tissue collection. For the fasting experiments, mice were deprived of food for 12, 24 or 30 h (8 AM to 2 PM next day) with free access to drinking water.

### Cell culture

*bmf^+/+^*, *bmf^+/-^* and *bmf^-/-^* mouse embryonic fibroblasts (MEFs) were isolated from embryonic day 14.5 embryos by trypsin digestion of the remaining tissue after removal of internal organs, brain, and fetal liver. MEFs were cultured in high-glucose version of Dulbecco’s modified Eagle’s medium (DMEM)(VWR Life Science) supplemented with L-glutamine (Gibco by Life Technologies), penicillin/streptomycin (Gibco by Life Technologies), 0.09 M of 2-mercaptoethanol (Sigma) and 10% FBS (Atlanta Biologicals). MEFs were genotyped as described previously (Labi et al., 2008). *Tsc2^+/+^* and *Tsc2^-/-^* mouse cells were provided by Elizabeth Henske (Brigham and Women’s Hospital, Boston, Massachusetts) and were cultured in high glucose DMEM supplemented with L-glutamine, penicillin/streptomycin, and 10% FBS. MAECs were prepared essentially as described previously (You, Richer et al., 2002) by incubation in pronase solution (1.4 mg/ml pronase and 0.1 mg/ml DNAse in DMEM) overnight at 4°C to dissociate airway epithelial cells from the basal lamina. Cells were collected by gently rocking the tracheas in DMEM/F-12 (Ham) media (Gibco by Life technologies) followed by centrifugation at 400 g for 10 min at 4°C. MAECs were grown in media described previously (Mou, Vinarsky et al., 2016). Two immortalized human airway epithelial cells, N1 and N3, were provided by S. Randell (University of North Carolina Chapel Hill, Chapel Hill, NC) (Fulcher, Gabriel et al., 2005). All airway epithelial cells were driven to differentiation by maintaining them on air-liquid interface culture conditions on Transwell membranes (Corning Incorporated COSTAR). AALEB cells are immortalized HAECs that have been well characterized (Lundberg, Randell et al., 2002). N1, N3 and AALEB cells were maintained in bronchial epithelial growth medium BEGM (Lonza) supplemented with growth factors (BEGM Singlequots, Lonza). Phoenix and HEK293T cells were cultured in high-glucose version of DMEM supplemented with L-glutamine, penicillin/streptomycin and 10 % FBS. Earle’s Balanced Salt Solution (EBSS) from Sigma was used as starvation media. The inhibitor pp242 (Sigma) was used at 0.5 or 2.5 µM to inhibit mTor kinase activity, MG132 (Sigma) was used at 10 µM to inhibit proteasomal degradation, protease inhibitor cocktail (Sigma) was used in all the tissues homogenized and cell lysates.

### Polymerase Chain Reaction

Total RNA from cultured cells was extracted by using TriReagent (Molecular Research Center, Inc.) according to the manufacturer’s instructions and RNA concentration was determined using a spectrophotometer Nano Drop 1000 (Thermo Scientific). The primer/probe sets for Bmf, CDKN1B, Muc5AC and p62 were obtained from Applied Biosystems. Target mRNAs were amplified by quantitative real time PCR in 20 µl reactions on the real-time PCR system 7900HT Sequence Detection System ABI PRISM (Applied Biosystems) using one step TaqMan RT enzyme and TaqMan RT-PCR Mix (Applied Biosystems). Relative quantification from triplicate amplifications were calculated by normalizing averaged threshold cycle (Ct) values to CDKN1B to obtain ΔCt, and the relative standard curve method was used for determining the fold change.

### Retroviral Silencing Using Short Hairpin RNA

Retroviral silencing vector encoding for human Bmf short hairpin RNA (shRNA) and control vector were used. Bmf-specific shRNA plasmids and control plasmids were purchased from OriGene (OriGene Technologies, Inc.) and were packaged into retroviral particles using Phoenix cells as specified by the manufacturer’s instructions. The harvested virus was then used to infect cells. To generate stable transfected cells, puromycin (Calbiochem) selection was used according to the manufacturer’s instructions. The shRNA knockdown efficiency was confirmed by quantitative RT-PCR.

### Cell Survival Assay

Cells were plated in 24-wells plate and when reaching 80-90% confluency were treated with the regular media (control), 0.5 µM pp242 in media, or EBSS (starvation). After 24 h the cells were washed, trypsinized and the viable cells were counted using Trypan Blue Solution (Sigma). The percent of cell survival was calculated relative to the surviving cells in the control media for each genotype.

### Adenoviral infection

Adenoviral expression vectors for GFP and Bmf_CUG_ were previously described (Contreras et al, 2013). Cells were plated, when reaching 80 % confluence they were infected with adenovirus with 50 or 100, MOI and 24 h later the infected cells were collected and analyzed.

### Transient Transfection

Transient transfections were carried out with the TransIT-2020 Reagent (Mirus) according to manufacturer’s protocol. Plasmid transfected were the following: pRK5-HA-UbiquitinK48 and pRK5-HA-UbiquitinK63 (Addgene plasmids # 17605 and #17606 respectively); prcCMV (used as an empty vector, E.v.), prcCMVbcl-2 (expressing wild type Bcl-2) and pcrCMVbcl2cb5 (expressing ER localized Bcl-2, ER-Bcl-2) were provided by Elizabeth Osterlund (Sunnybrook Research Institute, Toronto, ON, Canada), Bmf wild type (Bmf_S_ in pShuttle-CMV), BH3 mutant and DBD mutant. The Bmf mutant plasmids BH3 and DBD were generated by Mutagenex Inc. by 4 single-site substitutions to modify the DBD domain from DKATQTLS to DKAAAALS (DBD mutant) and by 3 single-site substitutions to modify the BH3 domain from IARKLQCIADQFHRL to IARKAQCIAAQFHRL (BH3 mutant).

### Western Blot Analysis

Fractions of cellular extracts proteins, quantified by Pierce BCA Protein Assay (Thermo Scientific) were mixed with 5x SDS sample buffer (312.5 mM Tris,10 % SDS, 5 % 2-mercaptoethanol, 0.05 % Bromophenol blue and 50 % glycerol; pH 6.8) and subjected to SDS-polyacrylamide gel electrophoresis using Tris/Glycine buffer system based on the method described by Laemmli (Laemmli, 1970). Precision Plus Protein Standards (BioRad) were used in every gel as molecular weight standards. After electrophoresis, proteins were transferred to a nitrocellulose transfer membrane (Bio-Rad). Membranes were blocked, incubated with primary antibody overnight at 4°C, washed, incubated with peroxidase-conjugated Affini Pure goat anti-rabbit or anti-rat, or rabbit anti-mouse or anti-goat secondary antibody (Jackson Immuno Research Laboratories, Inc.) for 1 hour, washed and visualized by Western Lightning Plus-ECL (Perkin Elmer) according to manufacturer’s protocol with the Luminiscent Image Analyzer LAS-4000 (Fujifilm). The following antibodies rabbit anti bak (#12105), rabbit anti bcl-2 (#3498 and #4223), rabbit anti Beclin1 (#3738), rabbit anti BiP (#3177), mouse anti CoxIV (#11967), rabbit anti cytochrome c (#11940), rabbit anti HA (#3724), rabbit anti LC3 (#2775), rabbit anti K48 ubiquitin (#8081), rabbit anti K63 ubiquitin (#5621), rabbit anti mitofusin2 (#9482), rabbit anti phospho-P70S6K (#9205), rabbit anti ubiquitin (#3933) were used at 1/1000 dilution and were obtained from Cell Signaling Technologies. Mouse anti-actin (A2228) is from Sigma and it was used at 1/4000 dilution. Mouse anti p62 (#610832) is from B.D. Transduction Labs and it was used in 1/1000 dilution. Rat anti Bmf (17A9) (ALX-804-508-C100) is from ENZO and it was used at 1/500 dilution. From Santa Cruz, we obtained rabbit anti DLC1/2 (sc-13969) used in 1/1000 dilution, mouse anti bcl-2 (C-2) (sc-7382) used in 1/300 dilution, mouse anti beclin1 (sc-48341) used in 1/500 dilution, mouse anti GAPDH (sc-32233) used in 1/1000 dilution and rabbit anti GFP (sc-8334) used in 1/500 dilution.

### Agarose Gels for Mucins separation and Western Blot Analysis

Mucin separation was performed in 0.8 % agarose gels as described previously (Ramsey, Rushton et al., 2016), and the transference to Nitrocellulose membranes was performed by capillarity. Membranes were blocked and incubated with primary and secondary antibodies as described in the previous section.

### Immunoprecipitation

Cells were lysed with IP Lysis/Wash buffer (0.025 M Tris, 0.15 M NaCl, 0.001 M EDTA, 1 % NP-40, 5 % glycerol; pH 7.4) and protein was quantified by BCA Protein Assay Pierce (Thermo Scientific). Between 300 and 500 µg of proteins were incubated with 1 µg of antibody (when using Santa Cruz or Abcam antibodies) or with the concentration recommended by the manufacturer (when using Cell Signaling antibodies) overnight at 4°C. Next day, 20 µl of Protein A agarose beads (Cell Signaling) were added and incubation continued for 3-5 hours at 4°C. Beads were washed with IP Lysis/Wash buffer 5 times and the proteins immunoprecipitated were collected by adding 5x SDS sample buffer. Inputs and immunopecipitations were analyzed by Western Blot. We immunoprecipitated proteins with mouse anti Bcl-2 (C-2) (sc-7382 from Santa Cruz), rabbit anti Beclin1 (ab55878 from Abcam) or mouse anti Beclin1 (sc-48341 from Santa Cruz).

### Histology and Morphometry

Mice were euthanized by lethal injection of euthasol, and lungs were inflated and fixed with Zn-formalin at a constant hydrostatic pressure of 25 cm for 2 h and further immersed in fixative for 48 h. The inflated lungs were embedded in paraffin and 5 µm sagittal sections were stained with Alcian Blue/Hematoxylin-Eosin. Morphometric analysis of alveolar volume measurement was carried out using the Visiomorph module of VisioPharm analysis software (Visiopharm).

### Immunofluorescence Assay

*bmf^-/-^* MEFs cells grown on 2 chamber slide (Lab-Tek II Chamber slide, Thermo Fisher Scientific) were transfected and 24 h later they were washed 3 times in PBS. The cells were fixed by treatment with 100% ice-cold methanol (which has been previously kept at −20°C for a minimum of 6 h) for 10 min on ice, according to a described protocol (Mebratu, Leyva-Baca et al., 2017). After fixation, the cells were washed 3 times for 5 min with 2% BSA in PBS and blocked for 1 h at RT in the same solution. Cells were incubated with rabbit anti LC3 antibodies (#2775 from Cell Signaling Technologies, at 1/800 dilution) for overnight at 4°C. After washing, cells were incubated with secondary antibody (Alexa Fluor 647 F(ab)’2 fragment goat anti-rabbit IgG (H + L) from Invitrogen by Thermo Fisher Scientific) for 2h at room temperature, washed and mounted with DAPI Fluoromount-G (Southern Biotech). Cells were viewed and images were taken by using the 63x oil DIC objective from a Leica DMi8 inverted confocal microscope and the LASX acquisition software. Images were processed by the Huygens deconvolution software and they were analyzed using the Slidebook 6 software.

*Bmf^+/+^*, *bmf^+/-^* and *bmf^-/-^* MAECs growing on Transwell membranes were washed, fixed with 4 % paraformadehyde for 30 min at room temperature, washed again and blocked for 1 hour at room temperature in blocking solution (3% BSA, 1% gelatin from cold water fish skin, 0.2% Triton X-100, 0.2% Saponin and 1% normal donkey serum in PBS). Cells were incubated with mouse anti Muc5AC (1/500 dilution) overnight in blocking solution at 4°C. After washing, cells were incubated with secondary antibody (Alexa Fluor 647 F(ab)’2 fragment goat anti-rabbit IgG (H + L) from Invitrogen by Thermo Fisher Scientific) for 2h at room temperature, washed and mounted with DAPI Fluoromount-G (Southern Biotech). Cells were viewed and images were taken by using Axioplan 2 (Carl Zeiss) with a Plan Aprochromal 63x/1.4 oil objective and a charge-coupled device camera (SensiCam; PCO), and the acquisition software used was Slide Book 6.0 (Intelligent Imaging Innovation).

### Statistical Analysis

Data are presented as means ± standard errors (SEM). Differences among groups were examined by one-way ANOVA when multiple groups were present or by t-test using Prism Statistical analysis software (GraphPad Software Inc.). A *P* value of <0.05 was considered significant.

## Author contributions

YT takes responsibility for the content of the manuscript, including the conception of the study. YT and MD made contributions to the design of the studies. YM, MD, SF, and DT carried out the described experiments, and all authors were involved with acquisition, analysis and interpretation of data and contributed to the writing of the manuscript.

## Acknowledgements

These studies were supported by grants from the National Institutes of Health (HL068111 and HL140839).

